# Bolt: A new age peptide search engine for comprehensive MS/MS sequencing through vast protein databases in minutes

**DOI:** 10.1101/551622

**Authors:** Amol Prakash, Shadab Ahmad, Swetaketu Majumder, Conor Jenkins, Ben Orsburn

## Abstract

The standard platform for proteomics experiments today is mass spectrometry, particularly for samples derived from complex matrices. Recent increases in mass spectrometry sequencing speed, sensitivity and resolution now permit comprehensive coverage of even the most precious and limited samples, particularly when coupled with improvements in protein extraction techniques and chromatographic separation.

However, the results obtained from laborious sample extraction and expensive instrumentation are often hindered by a sub optimal data processing pipelines. One critical data processing piece is peptide sequencing which is most commonly done through database search engines. In almost all MS/MS search engines users must limit their search space due to time constraints and q-value considerations. In nearly all experiments, the search is limited to a canonical database that typically does not reflect the individual genetic variations of the organism being studied. Searching for posttranslational modifications can exponentially increase the search space thus careful consideration must be used during the selection process. In addition, engines will nearly always assume the presence of only fully tryptic peptides. Despite these stringent parameters, proteomic data searches may take hours or even days to complete and opening even one of these criteria to more realistic biological settings will lead to detrimental increases in search time on expensive and custom data processing towers. Even on high performance servers, these search engines are computationally expensive, and most users decide to dial back their search parameters. We present Bolt, a new search engine that can search more than nine hundred thousand protein sequences (canonical, isoform, mutations, and contaminants) with 31 post translation modifications and N-terminal and C-terminal partial tryptic search in a matter of minutes on a standard configuration laptop. Along with increases in speed, Bolt provides an additional benefit of improvement in high confidence identifications, as demonstrated by manual validation of unique peptides identified by Bolt that were missed with parallel searching using standard engines. When in disagreement, 67% of peptides identified by Bolt may be manually validated by strong fragmentation patterns, compared to 14% of peptides uniquely identified by SEQUEST. Bolt represents, to the best of our knowledge, the first fully scalable, cloud based quantitative proteomic solution that can be operated within a user-friendly GUI interface. Data are available via ProteomeXchange with identifier PXD012700.

**Abstract Graphic:** 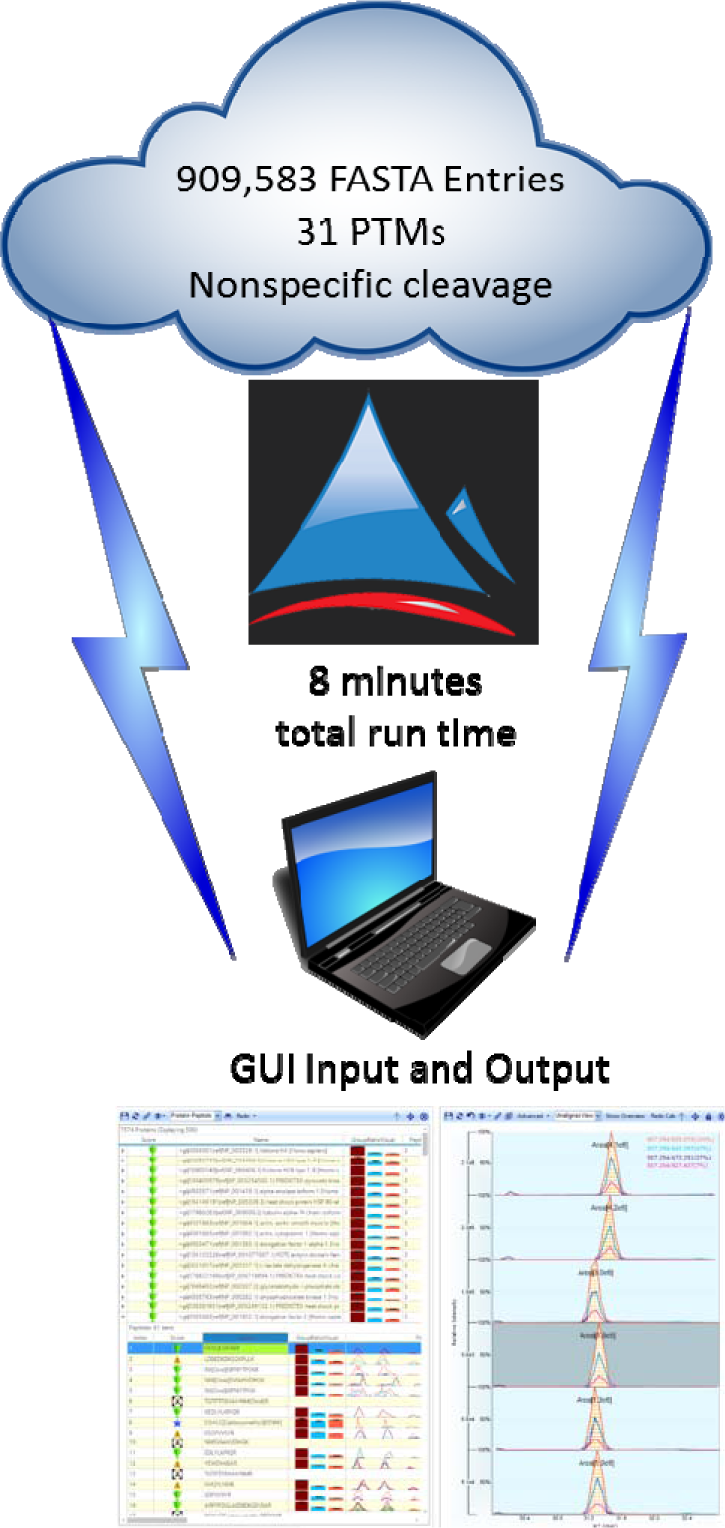

## Introduction

Mass spectrometry based proteomics studies are one of the most popular and powerful techniques for complex sample analysis. ^3^ With constant improvements in mass spectrometers speed and mass accuracy ^4^ and the recently proposed comprehensive data acquisition strategies like DIA ^5^, pSMART ^6^, and BOXCAR ^7^, it is possible to achieve a greater depth of proteome coverage today than what has been achievable in the past. The commonly used workflow involves extracting all the proteins from a sample (e.g., human serum or cell lysate), reducing/alkylating the cysteines, cleaving them with trypsin, and then loading the sample on a nanoflow HPLC coupled high resolution tandem mass spectrometer (LC-MS).^3^ After acquisition, the data is processed through a database search engine such as SEQUEST ^8^, Mascot ^9^ or Andromeda ^10^ that is nearly always housed on a powerful desktop computer. Samples are often precious and limited but still require comprehensive proteome coverage. Advances in sample preparation, chromatography and instrumentation have developed primarily to meet these needs at both large time and financial expense.^11^ The instruments cost hundreds of thousands of dollars to purchase and support. Furthermore, the overall workflow needs to be run by skilled professionals due to the complexity of the instrument operation and the lack of automated processes, presenting another costly aspect of the workflow.^12^ This intensive process is then concluded in most facilities by analysis with SEQUEST, a search engine that was designed for processing data from instruments that are no longer in use for proteomics applications ^13^. If users want their search algorithm to complete the analysis in a reasonable time (less than 30minutes), they have to select a canonical human database, select only a couple of most common post translation modifications (e.g., oxidation and N-term acetylation), and consider only fully tryptic peptides. The search algorithm’s restrictive design limits the protein search space and the ability for it to handle an excessive number of post translational modifications, thus forcing the user to make strategic trade-offs while using these algorithms to search their data. These hand-offs are not without biological consequence, as current generation instrumentation can easily identify post-translational modifications and non-tryptic cleavage events present in even the most complex human samples ^14^. Even if users had access to larger, pricy computational resources, they may still need to make limiting choices as otherwise the search engines might still take hours to process one analysis file.

Several recently developed search engines have tried to take aim at resolving some of issues that classical search engines possess. MSFragger ^15^and MetaMorpheus ^16,17^ can perform open searches, looking for an exceedingly large list of PTMs. While each of these newly built software suites provide an answer to one of the problems with classical search algorithms, they are still restricted by the overall size of the search space. They are unable to handle, in a timely fashion, the multi-dimensional increase in search space when important biological events such as protein splicing, isoforms, and mutations are added to the analysis.

In this paper, we introduce a modern search engine called Bolt that is capable of processing high resolution proteomics data against databases containing over nine hundred thousand protein entries in a matter of minutes. Canonical sequences, reviewed and non-reviewed isoforms, hundreds of thousands of known mutation variants, and a large contaminant database are all collected and searched using a semi-enzymatic digestion with a great number of PTMs, utilizing no greater computational resources for the end user than a standard laptop computer. Bolt consists of two main processing features, a client side algorithm that processes the analysis file and a high performance cloud server preconfigured to optimize the proteome search. We compare Bolt’s results to the most commonly used search engine, SEQUEST. Along with increased processing speed, Bolt also provides an additional benefit of high quality results, where 61% of the uniquely identified peptides by Bolt had strong fragmentation coverage compared to 10% for uniquely identified by SEQUEST.

## Material and Methods

Sample: All experiments were carried out with the HeLa digest standard and the Pierce Retention Time Calibration Standard (PRTC), both from Thermo Fisher. The PRTC was diluted with 0.1% formic acid to a total concentration of 50 fmol/µL. This solution of PRTC was used to reconstitute and dilute the HeLa digest standard. The HeLa standard was diluted to allow multiple injections ranging from 20ng to 500ng of peptide digest on column.

### LC/MS Platform

Two LC-MS systems were used for data acquisition as described:

An Orbitrap Fusion 1 system equipped with Tune version 3.0 installed in January 2018 with an EasyNLC 1200 system using 0.1% formic acid as Buffer A and 80% acetonitrile with 0.1% formic acid as buffer B. EasySpray 25cm columns with PepMap C-18 with 2um particle size and equivalent 2cm precolumn trap was used for all experiments described. Files from employing 2 distinct LC gradients were evaluated, with a total run time of 60 and 130 minutes, respectively. The gradients began a 5% B, ramping to 28% B within by 75% of their total acquisition time, to 50% B in the next 10%, followed by a rapid ramp to 98% B. Column equilibration to baseline conditions was performed automatically at the beginning of each run.

Additional files were acquired on a Q Exactive HF system using Exactive Tune 2.9 and equipped with MaxQuant Live 1.0.1. All source and chromatography conditions match that of the Fusion system as described above, with the exception that the system was an EasyNLC 1000. All components described are the products of Thermo Fisher Scientific. The mass spectrometry proteomics data have been deposited to the ProteomeXchange Consortium via the PRIDE^18^ partner repository with the dataset identifier PXD012700.

### SEQUEST/Proteome Discoverer

Proteome Discoverer 2.2 was used for the comparative searches, using the vendor provided default workflow for Q Exactive Basic Peptide ID. This consists of the SequestHT search engine with 10ppm MS1 tolerance, and 0.02 Da MS/MS tolerance. Dynamic modifications for iodoacetamide modification of cysteines and dynamic modifications were allowed for acetylation on the protein-N terminus, as well as for the oxidation of any methionine. Percolator was used for FDR according to manufacturer default settings. Up to 2 missed cleavages by trypsin were allowed. The files were searched against a UniProt/SwissProt FASTA downloaded in February 2018 that was parsed within the vendor software on the term “sapiens”. An in house generated contaminants database consisting of a combination of the MaxQuant common contaminants and the cRAP database (https://www.thegpm.org/crap/) was used to flag contaminants in both the processing and consensus workflows

### Bolt

The client-side processing (e.g., on the acquisition computer or user laptop) is used to read the vendor RAW file and extract the MS/MS spectra. The data is then sent for secondary processing to a high performance server (in our case Microsoft’s Azure Virtual Machine).The server has been pre-configured to a specific set of parameters that are commonly used for analysis of human proteomics samples. It currently holds a combination of human protein and contaminant sequences. As shown in Table 1, we used a combined database that contains 909,583 protein sequences. Approximately 8% of the protein sequences from the XMAn database are not considered in the current version of Bolt as those could not be mapped to the appropriate non-mutated protein sequence. We decided to use the entire Bovin database for this study of the cultured human HeLa cell line as we found more than 7,800 peptide sequences present in the Bovin database that are also present in the Trembl or SwissProt-Isoform database but not present in the SwissProt or Contaminant database. If we do not include the Bovin database, there is the risk of misidentifying a potential Bovin peptide as an isoform or a non-reviewed Trembl human peptide. The standard digestion configuration is for a tryptic digestion with two missed cleavage sites, partial tryptic search on both N- and C-terminus, Carboxymethyl or Carbamidomethyl as the alkylation agent along with the following 31 Post Translation Modifications: oxidation (M), phosphorylation (S), phosphorylation (T), phosphorylation (Y), methylation (K), methylation (R), pyroglutamate (N), pyroglutamate (Q), deamidation (N), deamidation (Q), hydroxyl (K), hydroxyl (P), formylation (S), formylation (T), formylation (K), formylation (N-term), dimethylation (K), dimethylation (R), dimethylation (N-term), trimethylation (K), trimethylation (R), acetylation (N-term), acetylation (K), carbamylation (K), carbamylation (C), carbamylation (R), carbamylation (n), propionylation (K), propionylation (N-term), GlyGly (K), lipoylation (K). Percolator is utilized to calculate a q-value using the target and decoy results. The Azure VM has 32 vCPUs to search in parallel. Once the high confidence target peptides are identified (user defined q-value <= 0.01), those are then transferred back to the client machine for further processing. Some rules are followed to reduce the search time and make the analysis easier. The number of PTMs was limited to 1 per peptide, but oxidation and N-terminal acetylation can still be considered as a second PTM. Peptides that have the same sequence but are different by only Leucine vs. Isoleucine are combined into single search. Aspartic acid is preferred over deamidation of asparagine if the rest of the sequence match is the same. Single PTM is preferred over two PTMs if there is no additional evidence; similarly, unmodified peptide is preferred over peptide with PTM if there is no additional evidence. Search tolerant is set as 10ppm for MS1 and 20ppm for MS/MS. Additionally, PTMs and partial cleavage peptides are allowed only for canonical database peptides.

**Table 1:** Various databases considered by Bolt to combine into a single sequence database.

| Protein database | Number of protein sequences |
| --- | --- |
| Human Swissprot (Canonical + Isoforms) | 42,439 |
| Bovin Swissprot (Canonical + Isoforms) | 23,882 |
| GPM CRAP | 116 |
| MaxQuant Contaminants | 245 |
| Human Trembl | 53,203 |
| XMAN Mutations | 872,125 |

### Pinnacle

In the current study, Bolt was run inside the Pinnacle software v 91 to help visualize and export results.

## Results

The Hela standard analyzed with a 130-minute gradient and a 60-minute gradient resulted in 53,921 and 41,384 MS/MS spectra respectively. Bolt was able to complete the reading of spectra, upload to server, complete search process, and send the data back to the local Pinnacle file system in 10.5 minutes and 8 minutes, respectively. The corresponding processing workflow run times for SEQUEST in Proteome Discover 2.3 were 48min and 37min. With the addition of the Proteome Discoverer consensus step, a total processing time reached above an hour for both cases. These run times do not include peptide quantitation times from either software as those are not relevant to this analysis.

Bolt reported 15,116 peptides and is compared to the 13,466 peptides identified by SEQUEST with q-values<=0.01 for the 130 minute standard gradient. Figure 2 shows the Venn diagram comparing these. 11,964 peptides were identified by both Bolt and SEQUEST, 2,870 peptides were uniquely identified by Bolt (Uniq-B), 959 were identified uniquely by SEQUEST (Uniq-S). The MS/MS spectra associated with the 282 peptides that were uniquely identified by Bolt had assignments to different peptides in the SEQUEST search (Mixed-B), correspondingly MS/MS spectra from 543 peptides that are uniquely identified by SEQUEST were assigned a different peptide by Bolt (Mixed-S). Next we wanted to assess whether the peptides identified uniquely by either software were correct. All the identified peptides were grouped into 4 classes: length greater than 12, 11 to 12, 9 to 10, and 7 to 8. Peptides with length 6 and smaller are not considered as those may yield even fewer fragments. Figure 2 plots the distribution of the number of matched fragment ions for each of these peptide length classes. For peptides longer than 12 residues, 85% of the peptides commonly identified by both search engines contained 8 or more fragment ions annotated in the MS/MS spectrum. Uniq-B has approximately 67% peptides with 8 or more fragment peak annotations. Comparatively, the Uniq-S group, only contained 14% peptides with 8 or more fragment peaks annotated. Common peptides, Uniq-B and Uniq-S peptides contained 6 or more annotated fragment peaks in the matched spectrum for 95%, 85% and 32% occurrences respectively. In addition, the Uniq-S group was comprised of 37% of the peptides only having 3 or fewer, annotated peaks in the MS/MS spectrum. The same trend was observed in the peptide lengths 11 to 12 (top right panel) and peptide lengths 9 to 10 (bottom left panel) where Common and Uniq-B had a higher propensity of peptides having more ions annotated in the MS/MS spectrum compared to Uniq-S. Trends for the groups Mixed-B vs. Mixed-S are similar to Uniq-B vs. Uniq-S, where Bolt is consistently reporting peptides with more ions explained compared to SEQUEST. For smaller length peptide group (length 7 to 8), most peptides from both search engines had fewer ions annotated in the MS/MS spectrum. Even here, Uniq-S also had a lower percentage (14%) for >=6 annotated ions for these smaller length peptides when compared to Common (36%) and Uniq-B (44%) groups. The number of peptides that belong in each group are provided in Supplementary Table 1.

On the Hela 60 minute gradient samples, Bolt reported 11,529 peptides and SEQUEST reported 10,843 peptides with q-values <= 0.01. Figure 1 right panel shows the Venn diagram for these two. 9,437 peptides were identified by both, 1,863 peptides were classified as Uniq-B and 984 were classified as Uniq-S. Besides these, 229 peptides were placed in the Mixed-B category and 422 peptides in Mixed-S. Figure 3 plots the distribution of the number of matched fragment ions for each of these peptide length groups. For peptides longer than 12 residues, the Common set contained approximately 74% peptides with 8 or more fragment ions annotated in the MS/MS spectrum, Uniq-B had 57% and Uniq-S had 11%. For Uniq-S, 75% peptides had five or less ions annotated for peptides longer than 12 residues, whereas this was observed for only 10% of the Common peptides and 24% of Uniq-B peptides. Similar trends were observed for other length peptides as well, where Uniq-S had significantly lower number of annotated ions compared to Common and Uniq-B. The number of peptides that belong in each group are provided in Supplementary Table 1. By opening the stringency of the q value cutoff to 0.05 in Bolt, 56% of the peptides in the hela-130 Uniq-S group and 47% of the peptides in the hela-60 Uniq-S group were identified by Bolt.

**Figure 1:**
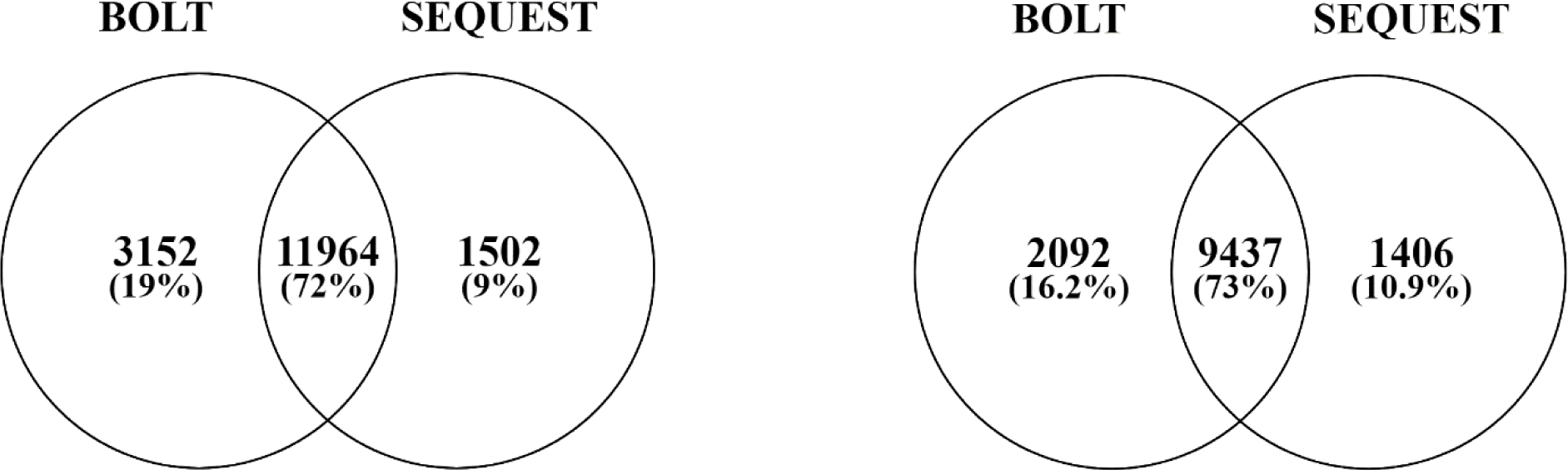
Venn diagram showing the overlap of peptide table from BOLT vs. SEQUEST. Left panel shows for hela-130 and right panel shows for hela-60.

**Figure 2:**
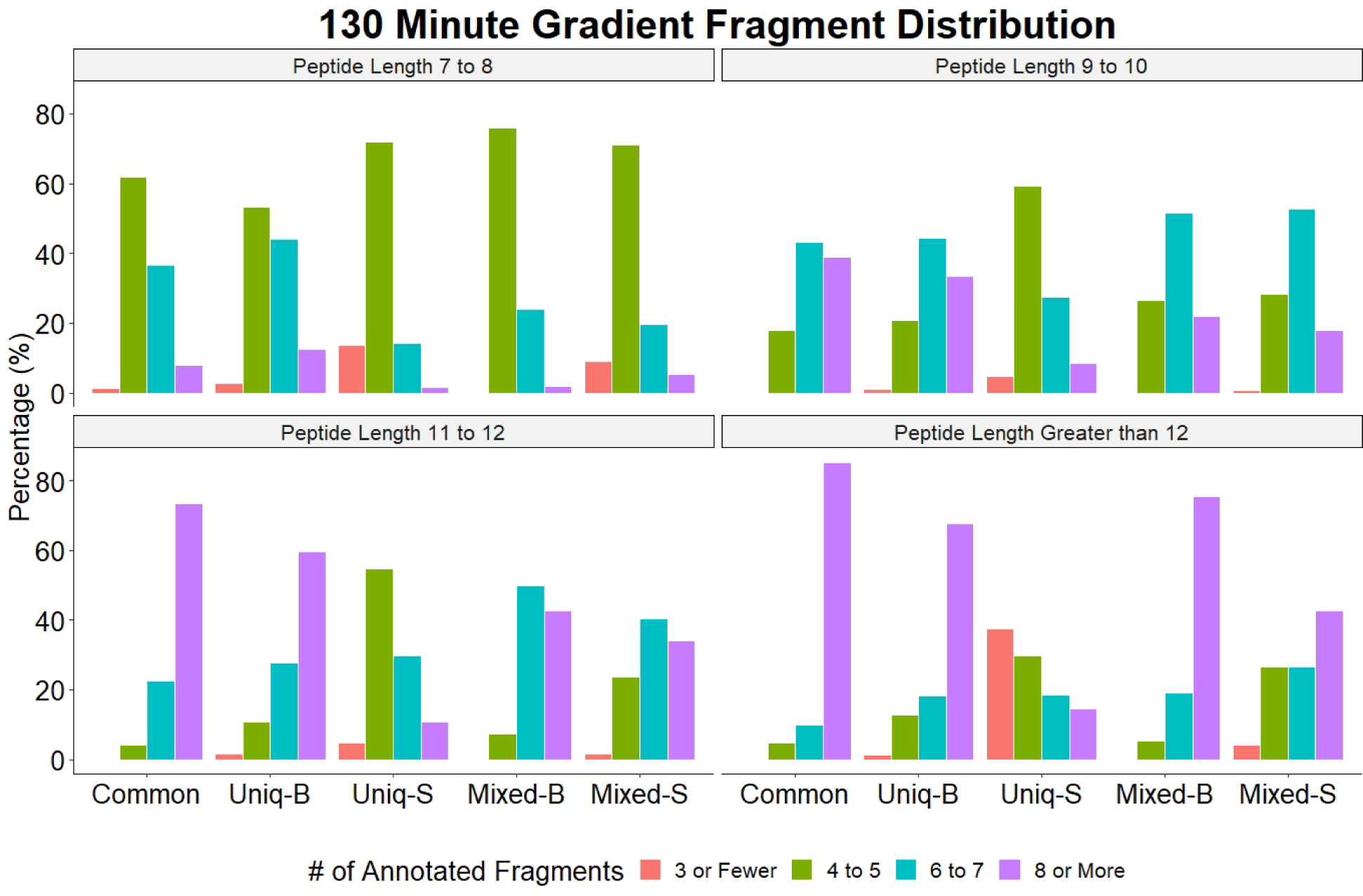
Distribution of number of assigned fragment ions in the spectrum matched to a peptide with qvalue<0.01 for hela-130 raw file. Top left panel shows the distribution for assigned ions for peptides that are longer than 12 amino acids. Distribution is shown for peptides that are both identified by Bolt and SEQUEST (‘Common’, shown in grey), identified uniquely by Bolt, identified uniquely by SEQUEST, peptides where Bolt generated a peptide ID from a spectrum that matched to something else by SEQUEST (Mixed-B), and peptides where SEQUEST generated a peptide ID from a spectrum that matched to something else by Bolt (Mixed-S). Matches by SEQUEST have significantly fewer ions explained compared to Bolt. Top right panel shows similar distributions for peptides length 11 and 12. Bottom left panel shows similar distribution for peptides length 9 and 10, and bottom right panel shows the same for peptides length 7 and 8.

**Figure 3:**
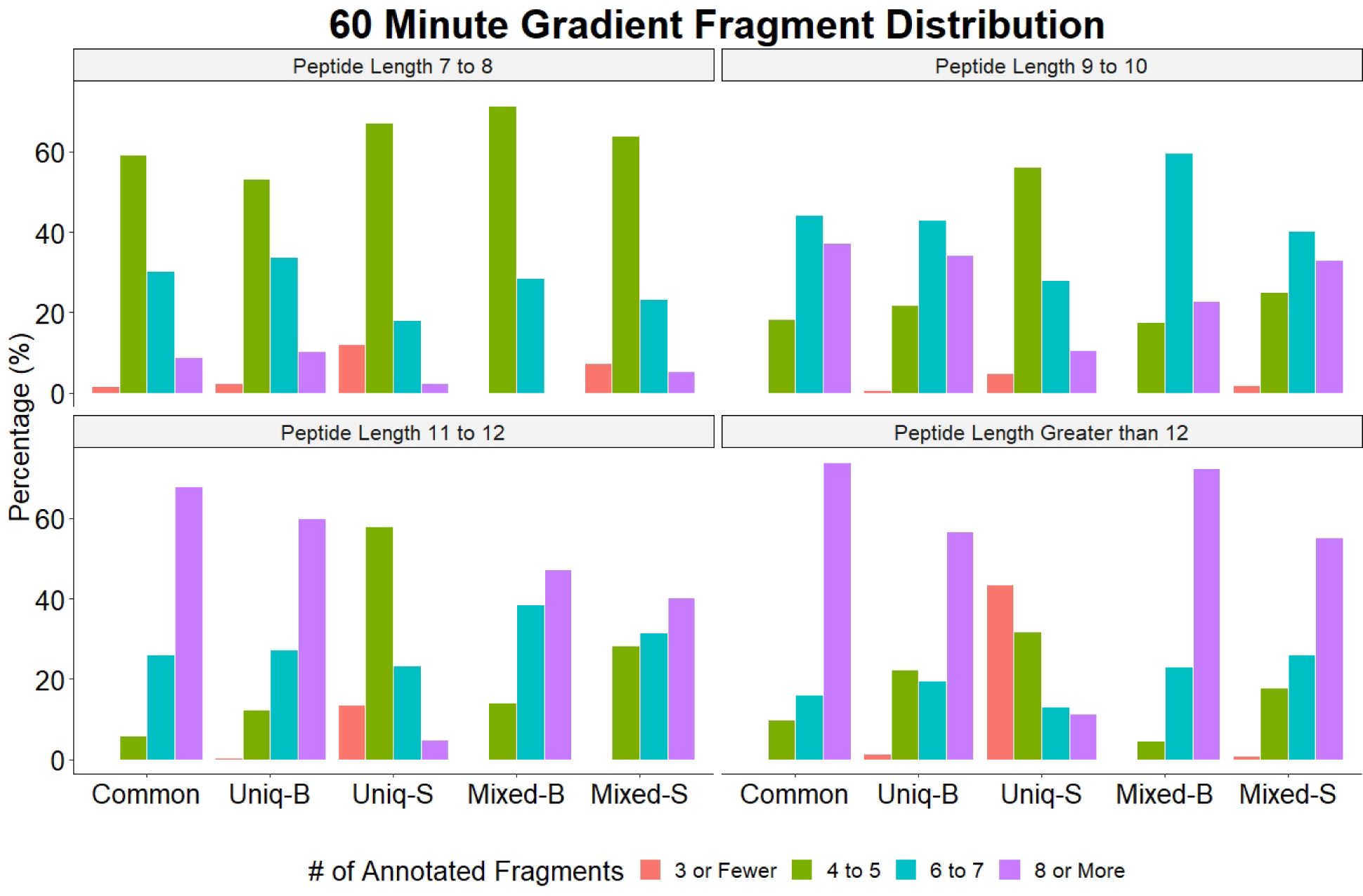
Distribution of number of assigned fragment ions in the spectrum matched to a peptide with qvalue<0.01 for hela-60 raw file. Top left panel shows the distribution for assigned ions for peptides that are longer than 12 amino acids. Distribution is shown for peptides that are both identified by Bolt and SEQUEST (‘Common’, shown in grey), identified uniquely by Bolt, identified uniquely by SEQUEST, peptides where Bolt generated a peptide ID from a spectrum that matched to something else by SEQUEST (Mixed-B), and peptides where SEQUEST generated a peptide ID from a spectrum that matched to something else by Bolt (Mixed-S). Matches by SEQUEST have significantly fewer ions explained compared to Bolt. Top right panel shows similar distributions for peptides length 11 and 12. Bottom left panel shows similar distribution for peptides length 9 and 10, and bottom right panel shows the same for peptides length 7 and 8.

One critical reason for running the different gradients was to observe how many of the peptide IDs from the shorter gradient can be observed in the longer gradient (thereby adding confidence in their assignment). We obviously do not expect to see 100% coverage of peptides from shorter gradient to longer gradient as even in otherwise identical experiment configurations. We expect about 70 to 80% reproducibility in the resulting peptide list. In the hela-60 raw file, Common group had 9,437 peptide IDs. From these 7,916 (83%) were identified in the hela-130 raw file (7,523 identified by both Bolt and SEQUEST, 232 identified by Bolt only and 160 identified by SEQUEST only). Uniq-S had 984 peptides out of which 574 (58%) peptides were identified in the hela-130 raw file (482 identified by both, 20 identified by Bolt only, 72 identified by SEQUEST only). Uniq-B had 1,863 peptides out of which 1,168 (62%) peptides were identified in the hela-130 raw file (337 identified by both, 817 identified by Bolt only, 14 identified by SEQUEST only). These results add even more confidence to the Uniq-B results.

One interesting aspect of the search engine is the capability to go through hundred thousands of published mutation sites in a matter of minutes. While both hela-60 and hela-130 reported 10 and 11 such mutated peptides found, two of these peptides were found common in both these analysis. For the protein GLRX3_HUMAN (O760003, Glutaredoxin-3) it is reported that residue 123 Proline is found in the mutated form as Serine. Thus the resulting tryptic peptide is HASSGSFLSSANEHLK (bold site shows site of mutation). Bolt found this peptide in both hela-60 and hela-130 raw files whereas SEQUEST did not assign this MS/MS a confident peptide ID. Figure 4 show the rich fragmentation patter in the hela-130 raw file, with most major ions assigned. The retention time observed in hela-60 is 11.7 minutes and in hela-130 is 12.1 minutes.

**Figure 4:**
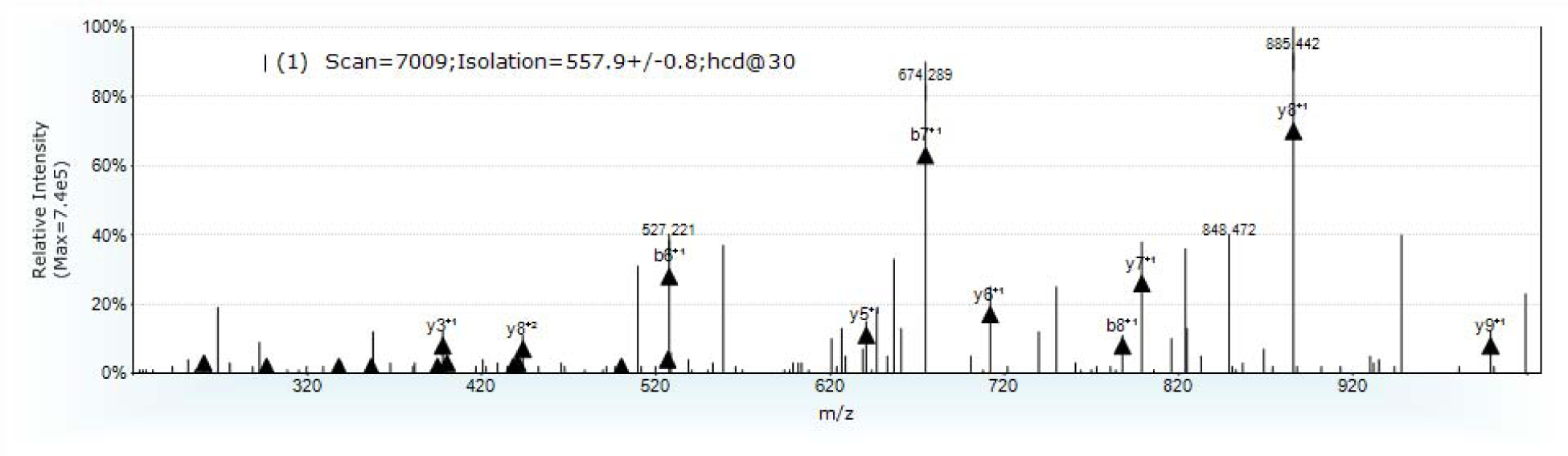
Fragmentation for the mutated peptide HASSGSFLSSANEHLK observed in hela-130 raw file as reported by Bolt.

Figure 5 shows the distribution of types of the Uniq-B group vs. the reported q-value. Bovin/Trembl/Mutation/Isoforms refer to peptides identified from the other sequence databases that were not provided to SEQUEST (Bovin database, TREMBL database, XMAn database, and SwissProt Isoform database respectively). These peptides are not present in the SwissProt Canonical database. Among the Bovin group peptides, 168 peptides (88%) from the hela-130 raw file and 149 peptides (92%) from the hela-60 raw file are present in the Contaminant database. While this may suggest that we do not need the entire Bovin database, as only 20-40 additional peptides from Bovin are found that do not exist in the common contaminant database, we have chosen here to err on the side of caution for a cell line grown in culture, rather than to report a Bovin peptide as a human peptide from the Trembl or XMAn mutation database. Alternatively, this also shows the need for a more complete contaminant database should be applied to current searches. The number of peptides for each type are provided in Supplementary Table 2.

**Figure 5:**
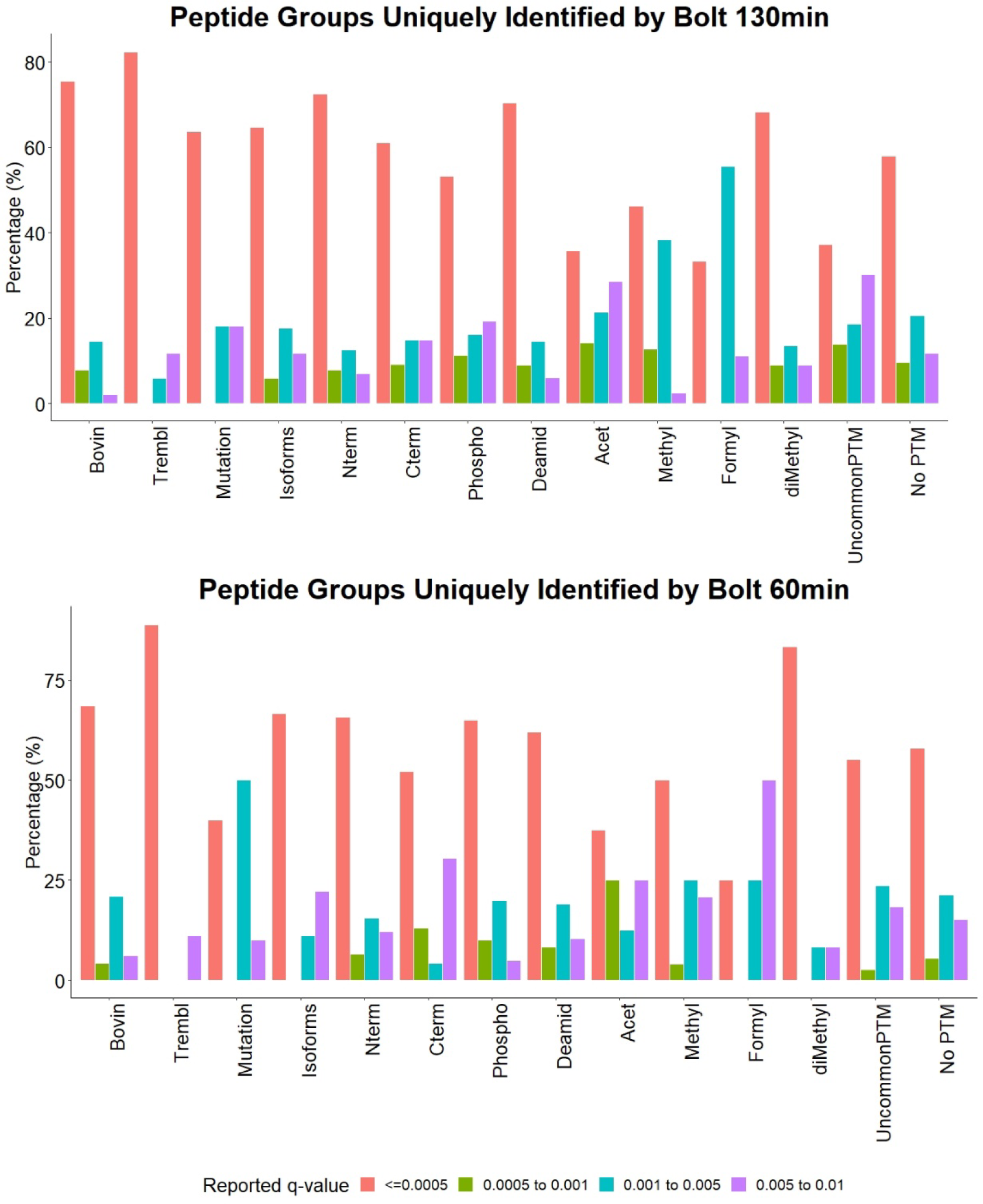
Distribution of types of peptides identified uniquely by Bolt (uniq-B group) vs. the reported q-value. Top panel shows results for the hela-130 raw file and the bottom panel shows for the hela-60 raw file. Bovin/Trembl/Mutation/Isoforms refer to peptides identified from the other sequence databases that were not provided to SEQUEST. Nterm/Cterm refer to peptides having tryptic cleavage site only on one terminal. Phospho/Deamid/Acet/Methyl refer to peptides having these PTMs. UncommonPTM refer to peptides having some uncommon PTMs that are considered by Bolt search. “No PTM” refers to tryptic peptides from the Swissport database that have no PTM other than oxidation.

In Figure 5, Nterm refers to SwissProt peptides having tryptic cleavage site only on the C-terminus, and Cterm refers to SwissProt peptides having a tryptic cleavage site only on the N-terminus. Almost 3% of the identified peptides are partially tryptic, thereby emphasizing the amount of information lost by ignoring partial tryptic search. The more commonly observed/studied PTMs are listed separately: Phospho/Deamid/Acet/Methyl refer to peptides having phosphorylation, deamidation or acetylation as a PTM. UncommonPTM refer to peptides having the uncommon PTMs that are considered by Bolt search (diMethylation, triMethylation, formylation, GlyGly, lipoylation, propionyl, carbamylation and hydroxylation). “No PTM” refers to tryptic peptides from the Swissport database that have no PTM other than oxidation. Figure 6 shows the distribution of number of annotated ions vs. peptide length for the Uniq-B-SwissProt group peptide so that it can be compared to the Uniq-S group distribution from Figures 2 and 3. For peptides longer than 12 residues, in the hela-130 raw file, Uniq-B-SwissProt had 77% peptides with at least 6 ions annotated whereas Uniq-S had 33% peptides. In the hela-60 raw file, for peptides longer than 12 residues, Uniq-B-SwissProt had 66% peptides with at least 6 ions annotated whereas Uniq-S had 25% peptides.

**Figure 6:**
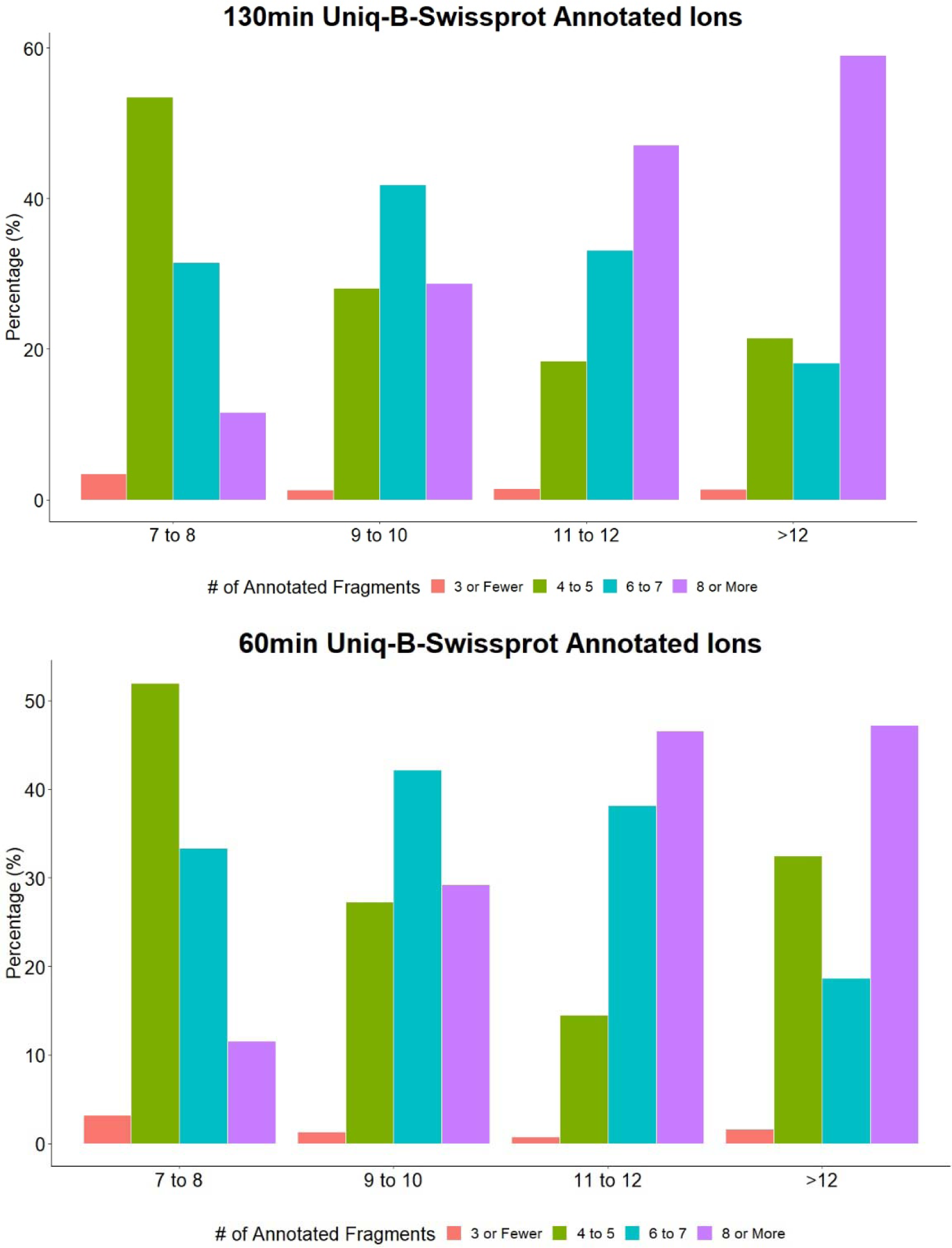
Distribution of number of annotated ions vs. peptide length for the Uniq-B-Swissprot group peptide (peptides uniquely identified by Bolt that belong to Swissprot database and do not have PTMs). Top panel is for the hela-130 run and bottom panel is for hela-60 run.

Investigating the Mixed-B and Mixed-S groups in more details, it appears these may arise due to chimeric spectra (i.e., a spectrum containing multiple peptide IDs). In these cases, the resulting sequence assignment from BOLT differs from that of SEQUEST. Evaluating them in more detail, we find that the top four annotated ions by one search engine are different from the top four annotated ions by the other search engine for more than 60% of such spectra. This further suggests that these could truly be chimeric spectrum where both search engines are reporting correct matches, but have different scoring and ranking algorithms. Figure 7 shows two such examples for hela-130 raw file. Top panel shows MS/MS spectrum #11131 assigned to peptide sequence VEEVGPYTYR by SEQUEST and to peptide SM[Oxid]QDVVEDFK by Bolt. Bottom panel shows MS/MS spectrum #8659 is assigned to peptide sequence VDSPTVTTTLK by SEQUEST and to peptide LGNTTVICGVK by Bolt. The marked annotated ions (almost complete y-ion series) are all different for both these assignments clearly indicating the presence of chimeric spectrum. All these peptides are from the SwissProt database, thus are present in the search space of both software.

**Figure 7:**
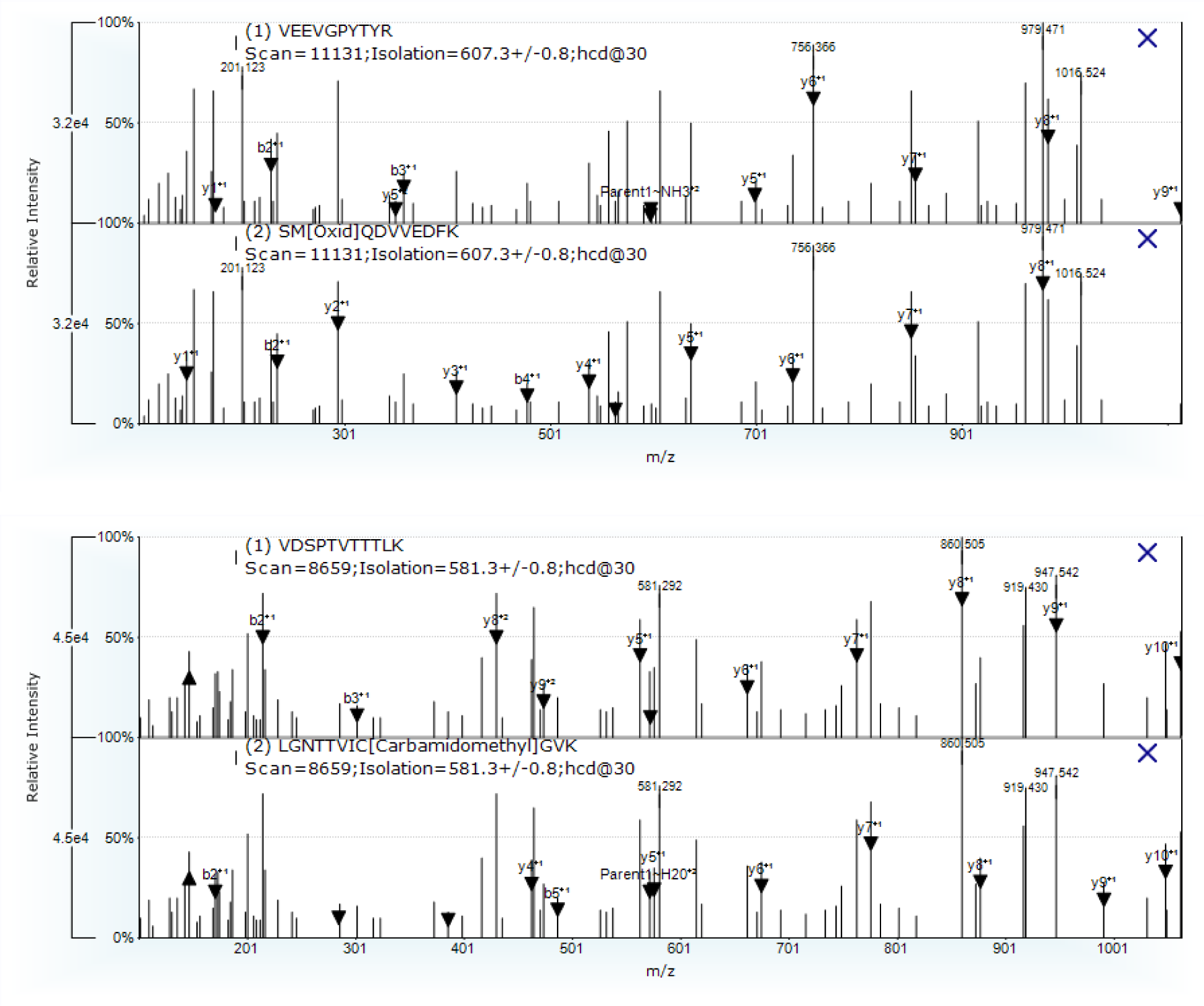
Two examples of chimeric spectrum in the hela-130 raw file. Top panel: MS/MS spectrum 11131 is assigned to peptide sequence VEEVGPYTYR by SEQUEST and to peptide SM[Oxid]QDVVEDFK by Bolt. Bottom panel: MS/MS spectrum 8659 is assigned to peptide sequence VDSPTVTTTLK by SEQUEST and to peptide LGNTTVICGVK by Bolt. Annotated ions are all different for both these assignments clearly indicating the presence of chimeric spectrum.

## Discussion

We have presented Bolt, a new search engine that is capable of searching over nine hundred thousand protein sequences, tens of dynamic PTMs, and partially cleaved peptides in a matter of minutes on a normal configuration computer. Selecting even one of these options in SEQUEST will make the processing take more than an hour on a high performance server and hours or even days on a more typical configuration computer^19^. If the goal of studying large proteomics cohorts is to sequence as deeply as possible, this makes Bolt a much more viable search engine than SEQUEST. For doing this search Bolt utilizes a high performance server on Azure cloud. The additional benefit of cloud server is that it can be scaled up or down as needed (e.g., more servers can be brought up in periods of high demand) with minimal cost association.

There are other search engines, e.g., Mascot, Byonic^20^, Andromeda, etc. that are used by many research groups. All of these have similar performance bottlenecks as SEQUEST - to have the search result be reported in a reasonable time, the user must limit the database and the list of dynamic post translation modifications as well as perform a search assuming near complete protease cleavage. A more extensive comparison with other search engines is in works and will be reported in a follow up study. Our expectation is that the unique and conflicting group of peptides will be different for each search engine but expect Bolt to still have a significant performance advantage over all other search engines.

While peptides with mutations may be the most interesting biologically, there are other classes of peptides that are also very important. Peptides from Trembl and Isoform databases help see protein variants that might have been ignored in previous studies. Peptides from Bovin can help improve the contaminant databases. Partially cleaved peptides can help identify protein fragments that are not present in the public databases, and these may be especially interesting. Peptides with other common and uncommon PTMs all help analyze various biological pathways. Thus, all these classes help enhance and complete the biological analysis in some form or the other. While we chose SwissProt and Trembl as the main databases, we can also choose ENSEMBL and RefSeq. We found both of these databases to have a large number of uncharacterized peptides (proteins) that are not part of our currently used databases. A recent study^21^ compared these various databases for proteogenomics but searching time and FDR were their primary bottlenecks. Bolt is a robust solution for both of these challenges, and plans to incorporate other proteomics databases (as well as variant databases) are in works. Our choice of PTMs for this study were derived from common human modifications that were of most relevant interest to other studies we currently have in works and served as a proof of principle We wanted to show that Bolt is capable of handling a large number of PTMs in an efficient manner without having to compromise on false discovery rate.

The results in Figure 2 and 3, demonstrate that identifications by Bolt are at least as good as SEQUEST and have fewer false positives in these samples. While we have used number of fragments as our primary representative measure, there are other criteria that one can use such as the percentage of top few MS/MS ions explained. However, most of these criteria are based on the assumption that the peptide produces a rich fragmentation pattern. For many classes of peptides this may not be true (e.g., peptides contain proline). We fully acknowledge that BOLT is still in early stages of development. Search engines like SEQUEST have been developed and optimized over many iterations and many years and have been optimized for downstream analysis with tools such as Percolator. It is quite possible that some of the identifications in the group Uniq-S are correct and Bolt is missing these. When we study some of these manually, some do appear to be correct. But without any further investigative studies and possibly synthesizing those peptides there is no way of knowing.

Another interesting area of study that we plan to investigate is chimeric spectrum. Figure 7 showed two examples of chimeric spectrum where both identifications by (Bolt and SEQUEST) seem correct. Some research groups have added capabilities to their search engines to process every spectrum as a possible chimeric spectrum^10,22^ and yield multiple identifications per MS/MS. However this often requires a second complete search and subsequently large increases in search time. Moreover, the comparison of results between search engines needs to be studied in greater depth when we allow multiple matches per MS/MS.

Bolt provides a truly modern option for setting up proteomics searches. In a very reasonable time, it can search through non-reviewed and mutation databases without loss of sensitivity. While we have shown and compared results on a peptide level, Bolt produces a protein list that is also longer than the one from SEQUEST’s output. This is expected, but it requires a careful evaluation of protein grouping algorithms for conclusive analysis, especially when there is a high degree of homology in the database. Bolt represents, to the best of our knowledge, the first fully scalable, cloud based quantitative proteomic solution that can be operated within a user-friendly GUI interface.

